# A CRISPRi-dCas9 system for archaea and its use to examine gene function during nitrogen fixation by *Methanosarcina acetivorans*

**DOI:** 10.1101/2020.06.15.153874

**Authors:** Ahmed E. Dhamad, Daniel J. Lessner

## Abstract

CRISPR-based systems are emerging as the premier method to manipulate many cellular processes. In this study, a simple and efficient CRISPR interference (CRISPRi) system for targeted gene repression in archaea was developed. The *Methanosarcina acetivorans* CRISPR-Cas9 system was repurposed by replacing Cas9 with the catalytically dead Cas9 (dCas9) to generate a CRISPRi-dCas9 system for targeted gene repression. To test the utility of the system, genes involved in nitrogen (N_2_) fixation were targeted for dCas9-mediated repression. First, the *nif* operon (*nifHI_1_I_2_DKEN*) that encodes molybdenum nitrogenase was targeted by separate guide RNAs (gRNA), one targeting the promoter and the other *nifD*. Remarkably, growth of *M. acetivorans* with N_2_ was abolished by dCas9-mediated repression of the *nif* operon with each gRNA. The abundance of *nif* transcripts was >90% reduced in both strains expressing the gRNAs, and NifD was not detected in cell lysate. Next, we targeted NifB, which is required for nitrogenase cofactor biogenesis. Expression of a gRNA targeting the coding sequence of NifB decreased *nifB* transcript abundance >85% and impaired but did not abolish growth of *M. acetivorans* with N_2_. Finally, to ascertain the ability to study gene regulation using CRISPRi-dCas9, *nrpR1* encoding a subunit of the repressor of the *nif* operon was targeted. The *nrpR1* repression strain grew normally with N_2_ but had increased *nif* operon transcript abundance consistent with a NrpR1 as repressor. These results highlight the utility of the system, whereby a single gRNA when expressed with dCas9 can block transcription of targeted genes and operons in *M. acetivorans*.

**IMPORTANCE:** Genetic tools are needed to understand and manipulate the biology of archaea, which serve critical roles in the biosphere. Methanogenic archaea (methanogens) are essential for the biological production of methane, an intermediate in the global carbon cycle, an important greenhouse gas and a biofuel. The CRISPRi-dCas9 system in the model methanogen *M. acetivorans* is the first Cas9-based CRISPR interference system in archaea. Results demonstrate that the system is remarkably efficient in targeted gene repression and provide new insight into nitrogen fixation by methanogens, the only archaea with nitrogenase. Overall, the CRISPRi-dCas9 system provides a simple, yet powerful, genetic tool to control the expression of target genes and operons in methanogens.

## INTRODUCTION

The CRISPR-Cas (<u>c</u>lustered <u>r</u>egularly <u>i</u>nterspaced <u>s</u>hort <u>p</u>alindromic <u>r</u>epeats-<u>C</u>RISPR <u>a</u>ssociated) system is a primitive adaptive immune system found in prokaryotes (bacteria and archaea) [1, 2]. Approximately 40% of bacteria and 90% of archaea contain a CRISPR-Cas system [3]. The system is used to combat foreign nucleic acid (e.g. virus) by using previously acquired nucleic acid to generate a guide RNA (gRNA) that targets the foreign nucleic acid for cleavage by a Cas endonuclease (e.g. Cas9). Several types of CRISPR systems have been discovered, and detailed characterization has led to the development of numerous genetic tools [4–6]. The type II CRISPR-Cas9 system has garnered the most applications, in part due to DNA targeting by Cas9 being extremely precise with virtually no off-target cleavage [7]. For example, systems have been designed to 1) edit genomes by using gRNA-directed Cas9-mediated DNA double-stranded breaks to allow gene deletion and additions using homology-dependent repair [8, 9], and 2) control gene expression using a nuclease-deficient or dead Cas9 (dCas9) protein that targets specific DNA but does not cut it [10]. DNA binding by dCas9 blocks transcription initiation and/or elongation of gRNA-targeted genes. The latter has been termed CRISPR interference (CRISPRi) [11]. Gene repression by CRISPRi-dCas9 systems is a powerful gene regulatory tool, whereby the expression of multiple gRNAs can allow the repression of several genes at the same time [10]. CRISPRi-dCas9 systems have been developed to control gene expression in bacteria (e.g. *E. coli*) and eukaryotes (e.g. yeast and mammalian cells) but not archaea [11]. Archaea play central roles in the cycling of key elements (e.gs. carbon, nitrogen, and iron) on Earth, and they serve as important sources for the development of biotechnology, including the production of biofuels and enzymes [12, 13]. CRISPRi systems based on endogenous type I and type III CRISPR systems have been developed for the extremophilic archaea *Haloferax volcanii* and *Sulflobus* spp., respectively. The *Sulfolobus* CRISPRi system is a post-transcriptional gene expression control system that utilizes a CRISPR type III complex to target mRNA for degradation [14–17]. The *Haloferax* CRISPRi system utilizes mutant strains deleted of part or all endogenous CRISPR loci and expression of gRNAs that direct the Cascade complex to block transcription of a targeted gene or operon [18, 19].

The strictly anaerobic methane-producing archaea (methanogens) are particularly important archaea. Methanogen-produced methane is critical to the global carbon cycle, is a greenhouse gas, and is a biofuel [13]. Methanogens are also the only archaea that are capable of dinitrogen (N_2_) fixation (diazotrophy) [20–22]. Methanogens were likely the first diazotrophs and the source of nitrogenase, the unique enzyme that catalyzes the energy intensive reduction of N_2_ into usable ammonia (NH_3_) [23]. Advanced genetic tools are necessary to understand the fundamental biology of methanogens and to provide platforms for genetic engineering to facilitate their use in biotechnology. Recently, a CRISPR-Cas9 system was developed to edit the genome of the model methanogen *Methanosarcina acetivorans* [24]. Genome editing by the CRISPR-Cas9 system is driven by native homology-dependent repair (HDR) or by coexpression of the nonhomologous end-joining repair (NHEJ) pathway. Cas9-mediated genome editing has increased the efficiency and speed of genetic analyses of *M. acetivorans*. However, mutant generation typically requires 4-5 weeks (including plasmid construction) and generating strains with multiple mutations can require sequential strain construction [24].

We sought to repurpose the *M. acetivorans* CRISPR-Cas9 system by replacing Cas9 with dCas9 to generate a system to control gene expression (i.e. CRISPRi-dCas9). To test the CRISPRi-dCas9 system, we selected target genes that are not essential, but when repressed would give an observable phenotype. *M. acetivorans* contains the *nif* operon that encodes the molybdenum (Mo) nitrogenase, as well as operons encoding the alternative vanadium (V) and iron (Fe) nitrogenases [25]. Due to substantial ATP consumption required to fix N_2_, nitrogenase production is highly regulated and is produced only in the absence of a usable nitrogen source (e.g. NH_3_) [26]. All diazotrophs contain Mo-nitrogenase, which uses the least amount of ATP [20]. The alternative nitrogenases are typically only used when molybdenum is not available [27]. Production of functional Mo-nitrogenase also requires unique cofactor biogenesis machinery [28]. Although *M. acetivorans* Mo-nitrogenase production and nitrogen fixation has not been investigated in detail, it provides a suitable target for the CRISPRi-dCas9 system since it is only required for growth in the absence of a fixed nitrogen source. Genes encoding the structural, regulatory, and cofactor biogenesis components of Mo-nitrogenase were targeted for dCas9-mediated repression. Results demonstrate that the CRISPRi-dCas9 system is remarkably efficient in target gene repression. The *M. acetivorans* CRISPRi-dCas9 system expands the archaeal genetic toolbox and will specifically expedite the investigation and genetic engineering of methanogens.

## RESULTS

### Design of a CRISPRi-dCas9 system in *M. acetivorans*

The CRISPR-Cas9 system developed by Nayak and Metcalf [24] was used as the foundation for the development of a *M. acetivorans* CRISPRi-dCas9 system. In the CRISPR-Cas9 system, one or more gRNAs are expressed from a methanol-inducible promoter (*P_MeOH_*) and Cas9 is expressed from a tetracycline-inducible promoter (*P_Tet_*). Specialized strains (e.g. WWM73) of *M. acetivorans* are transformed with a plasmid (e.g. pDN203) containing the gRNA(s) and Cas9. Plasmid pDN203 and derivatives can integrate into the chromosome of transformed cells by site-specific recombination or alternatively, can be converted into a replicating plasmid by co-integration with pAMG40 [24]. To convert the CRISPR-Cas9 system into a CRISPRi-dCas9 system, *cas9* in pDN203 was replaced with *dcas9* from pMJ841 to generate pDL730. Plasmid pDL730 lacks a gRNA and therefore serves as the universal plasmid to incorporate gRNAs. *M. acetivorans* containing pDL730 was the control strain used in all experiments.

CRISPRi-dCas9 strains of *M. acetivorans* were generated by integration of pDL730 and derivatives into the chromosome of parent strain WWM73. Alternatively, pDL730 could be converted into a replicating plasmid. In all CRISPRi-dCas9 strains, the repression of a target gene is induced by growth with methanol and the addition of tetracycline (**Fig. 1**). To test the ability of the system to silence or “knockdown” transcription of gRNA-targeted genes, we selected the *nif* operon, and related genes, for dCas9-mediated repression. These genes should only be expressed when *M. acetivorans* is not given a usable nitrogen source (e.g. NH_4_Cl).

**Fig. 1.**
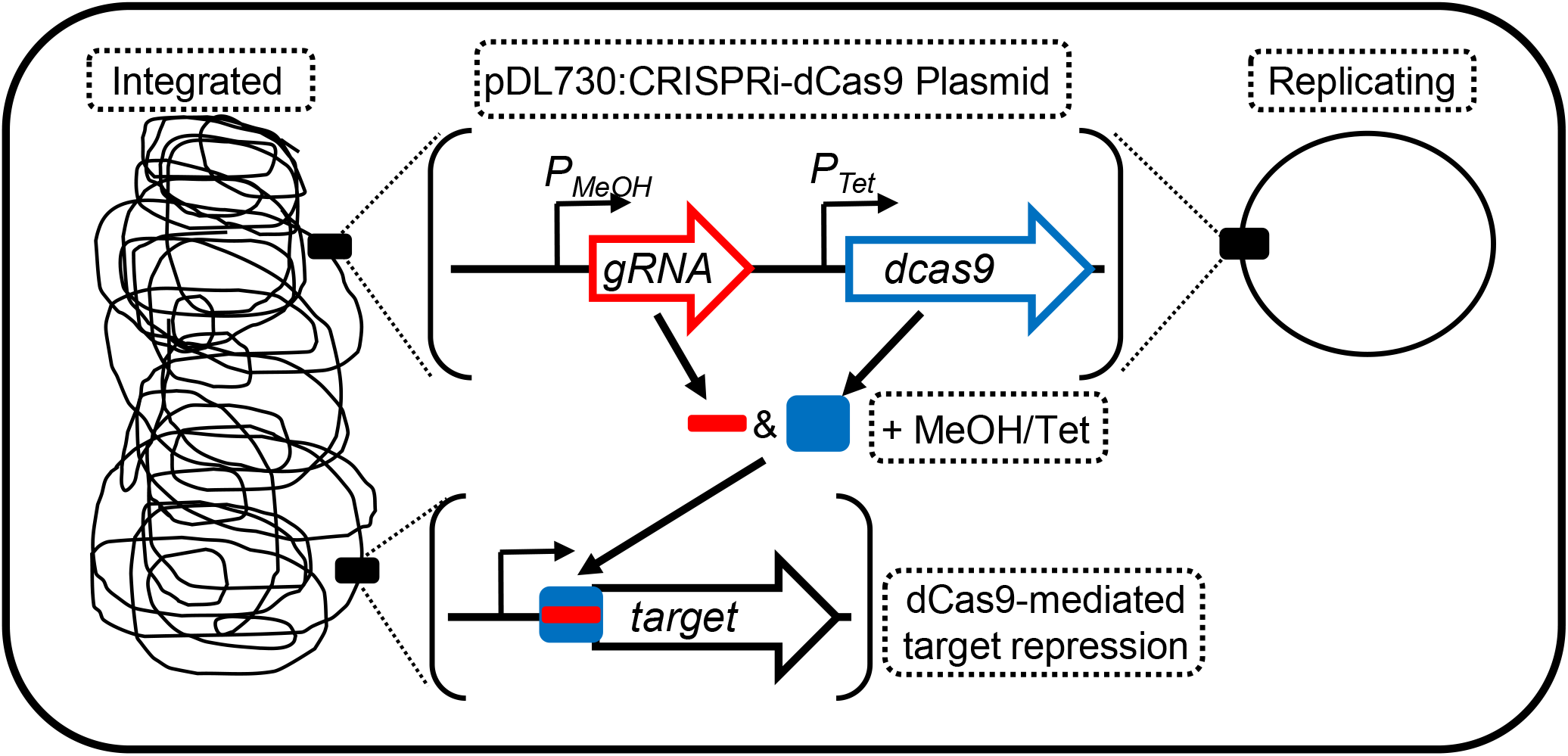
Design of a CRISPRi-dCas9 system for targeted gene repression in *M. acetivorans*. The CRISPRi-dCas9 plasmid pDL730 and derivatives can either integrate into the chromosome of *M. acetivorans* or be converted into a replicating plasmid. Expression of the gRNA and dCas9 are induced by growth with methanol (MeOH) and the addition of tetracycline (Tet).

However, expression of the *nif* operon in *M. acetivorans* has not been investigated. Therefore, diazotrophic growth and expression of the *nif* operon in response to available nitrogen was first determined.

### *M. acetivorans* expresses Mo-nitrogenase (NifHDK) and fixes N_2_ when grown in the absence of NH_4_Cl

To provide a baseline for CRISPRi-dCas9 experiments, the expression of Mo-nitrogenase and diazotrophic growth of *M. acetivorans* strain WWM73 was examined. Growth of strain WWM73 in sealed tubes with a headspace containing 75% N_2_ was compared in standard high-salt (HS) medium +/− NH_4_Cl, the preferred nitrogen source (**Fig. 2A**). The cell yield and growth rate of cultures lacking NH_4_Cl were lower compared to cultures containing NH_4_Cl, consistent with increased energy consumption needed to fix N_2_. HS medium also contains cysteine, a possible fixed nitrogen source. Recent evidence indicated *M. acetivorans* can use cysteine as a sole nitrogen source [29]. However, extensive experimentation with strain WWM73 and derivatives clearly demonstrates the inability of *M. acetivorans* to use cysteine as a nitrogen source (see below). Also, *M. acetivorans* is incapable of growing in cysteine-containing HS medium in tubes with a headspace devoid of N_2_ (data not shown). Moreover, Real-time quantitative PCR (qPCR) analyses revealed increased transcript abundance of both *nifH* and *nifD* in cells grown in medium lacking NH_4_Cl and containing cysteine (**Fig. 2B**). Finally, NifD was only detected by Western blot in lysate from cells grown in the absence of NH_4_Cl (**Fig. 2C**). These results demonstrate that expression of the *nif* operon is tightly controlled and production of Mo-nitrogenase occurs only in the absence of a usable nitrogen source (e.g. NH_4_Cl).

**Fig. 2.**
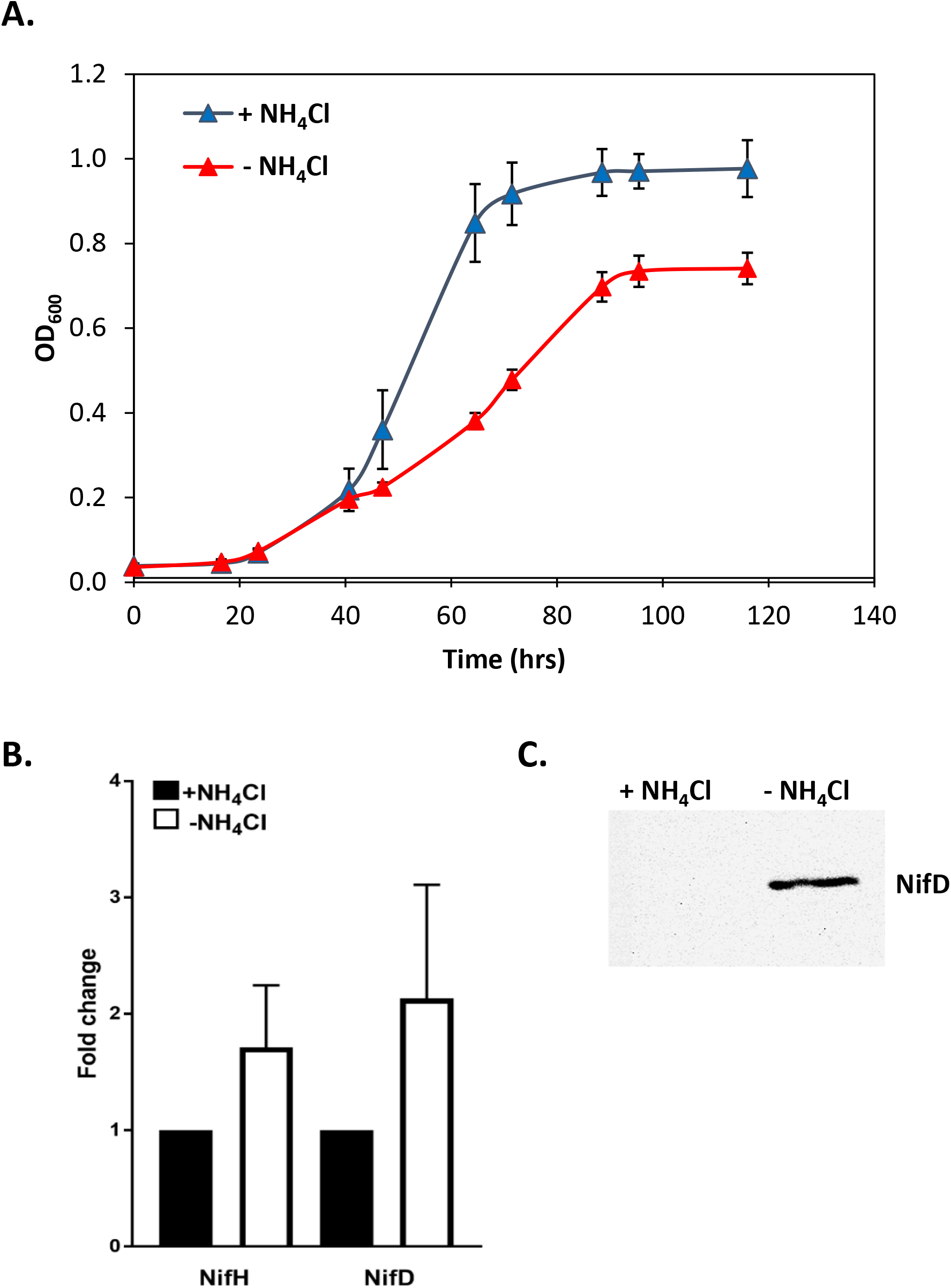
Effect of NH_4_Cl on *M. acetivorans* growth and *nif* operon expression. **A)** Growth in HS medium +/− NH_4_Cl with a 75% N_2_ headspace. **B**) Fold change in *nifH* and *nifD* transcript abundance in cells grown in medium +/− NH_4_Cl as determined by qPCR. **C)** Western blot analysis using NifD-specific antibodies of lysate from *M. acetivorans* cells grown +/− NH_4_Cl.

### Diazotrophic growth of *M. acetivorans* is abolished by dCas9-mediated repression of the *nif* operon

To test the ability of the CRISPRi-dCas9 system to repress expression of Mo-nitrogenase, gRNAs were designed to target two separate regions of the *nif* operon (**Fig. 3A**). NifHDK are the structural components of Mo-nitrogenase, NifI_1_I_2_ regulate nitrogenase activity, and NifEN serve as the M-cluster assembly scaffold [22]. NifHDK and NifEN are essential for N_2_ fixation by Mo-nitrogenase [23, 28]. A 20-nt gRNA (gRNA-*P_nif_*) was designed to target the sense (non-template) strand starting 15 nucleotides downstream of the transcription start site of the *nif* operon. A second gRNA (gRNA-*nifD*) targets the sense strand starting 100 nucleotides downstream of the start codon of *nifD*. Each gRNA was separately added to pDL730 to generate pDL731 and pDL732 (**Table S3**). *M. acetivorans* strain WWM73 was separately transformed with pDL731 or pDL732 to generate strains DJL74 and DJL76, respectively. Control strain DJL72 contains gRNA-free pDL730 (**Table 1**). To test the effect of each gRNA on diazotrophic growth of *M. acetivorans*, each strain was grown with methanol in HS medium +/− NH_4_Cl. Since dCas9 is expressed from a tetracycline-inducible promoter, the effect of the presence and absence of tetracycline on growth was also examined.

**Fig. 3.**
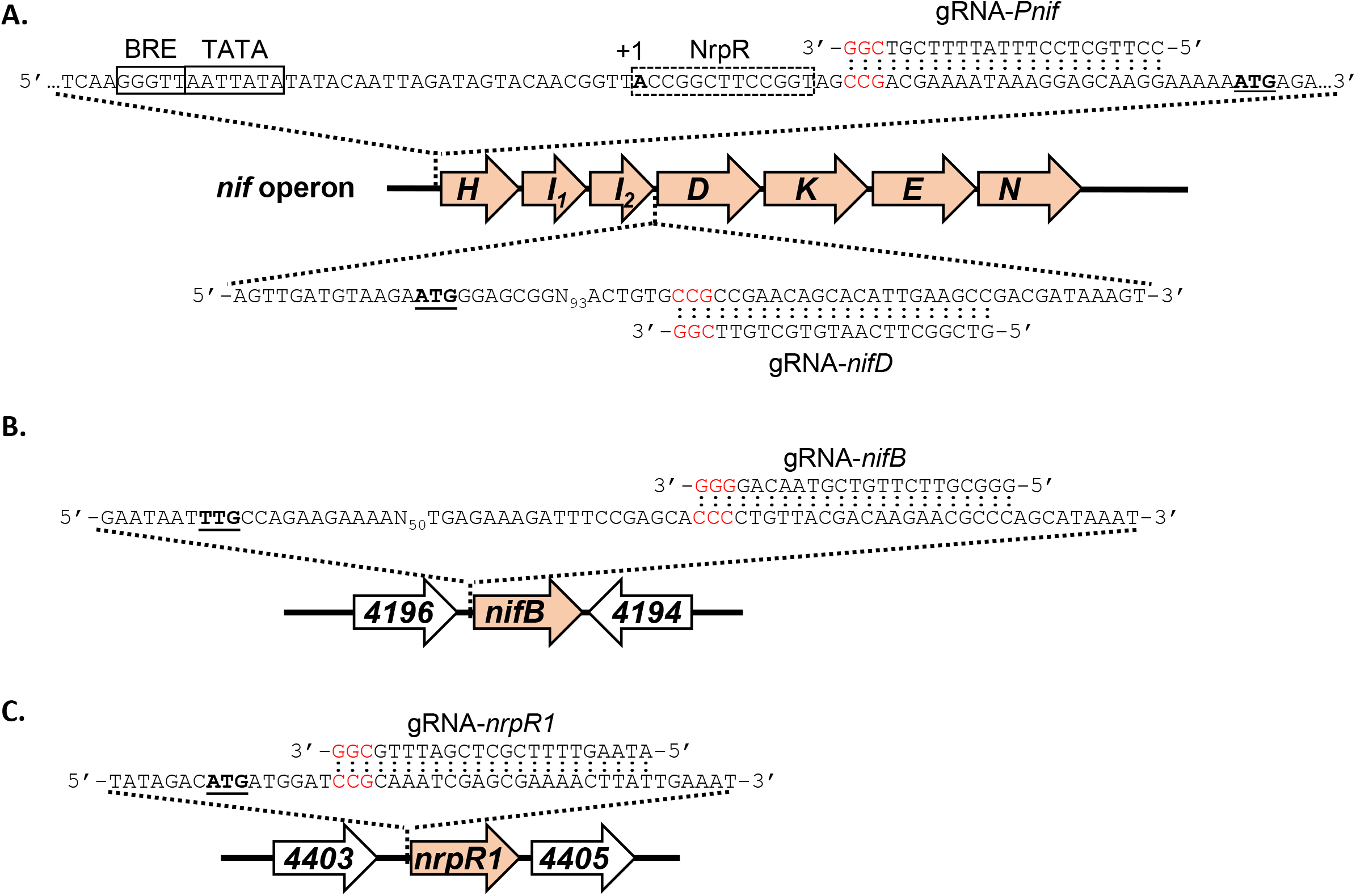
Designed gRNAs for targeted dCas9-mediated gene repression. **A)** gRNAs targeting the promoter (gRNA-*Pnif*) and 5’-end of *nifD* (gRNA-*nifD*) of the *nif* operon. The BRE, TATA, and NrpR operator sequences are boxed**. B)** gRNA targeting the 5’-end of *nifB.* **C)** gRNA targeting the 5’-end of *nrpR1*. Start codons are bold and underlined. PAM sequences are red.

**Table 1.** *M. acetivorans* strains used in this study.

| Strain | Genotype | gRNA | Source |
| --- | --- | --- | --- |
| WWM73 | Δhpt::PmcrB-tetR-ϕC31-int-attP | none | [46] |
| DJL72 | WWM73 att::pDL730 | none | This study |
| DJL74 | WWM73 att::pDL732 | gRNA-*Pnif* | This study |
| DJL76 | WWM73 att::pDL731 | gRNA-*nifD* | This study |
| DJL79 | WWM73 att::pDL739 | gRNA-*nrpR1* | This study |
| DJL105 | WWM73 att::pDL745 | gRNA-*nifB* | This study |

*M. acetivorans* strain DJL74 grew identical to strain DJL72 in medium with NH_4_Cl with or without tetracycline (**Fig. 4A**), indicating expression of gRNA-*P_nif_* does not impact growth with NH_4_Cl. However, whereas strain DJL72 was able to grow by fixing N_2_ in medium lacking NH_4_Cl, diazotrophic growth of strain DJL74 was abolished (**Fig. 4A**). Strain DJL74 was only able to grow using initial NH_4_Cl carried over from the inoculum to reach an OD_600_ of ~ 0.2 but was unable to switch to diazotrophic growth. The growth profiles of strain DJL74 were identical in the presence and absence of tetracycline, indicating that sufficient dCas9 is produced even in the absence of the inducer. Remarkably, the cessation of growth was absolute; the cells were unable to overcome dCas9-mediated repression of the *nif* operon as the OD_600_ stayed at ~0.2 for several days. To confirm that the inability of strain DJL74 to grow was due to the lack of fixed nitrogen as a result of dCas9-mediated repression of the *nif* operon, NH_4_Cl was added to some of the culture tubes ~100 hours after the cessation of growth. Cells in these cultures immediately started to grow and reached a final OD_600_ comparable to cultures grown initially with NH_4_Cl (**Fig. 4A**). A *M. acetivorans* strain with the replicating version of pDL732 exhibited the same phenotype (**Fig. S1**), indicating the location of the CRISPRi-dCas9 plasmid does not impact system function. Expression of gRNA-*nifD* also abolished diazotrophic growth of strain DJL76 in the absence of NH_4_Cl (**Fig. 4B**). The growth profiles of the strains expressing gRNA-*Pnif* and gRNA-*nifD* were identical (**Fig. 4)**. Thus, targeting either the *nif* operon promoter or downstream of the start codon of *nifD* results in dCas9-mediated repression sufficient to abolish diazotrophic growth of *M. acetivorans*.

**Fig. 4.**
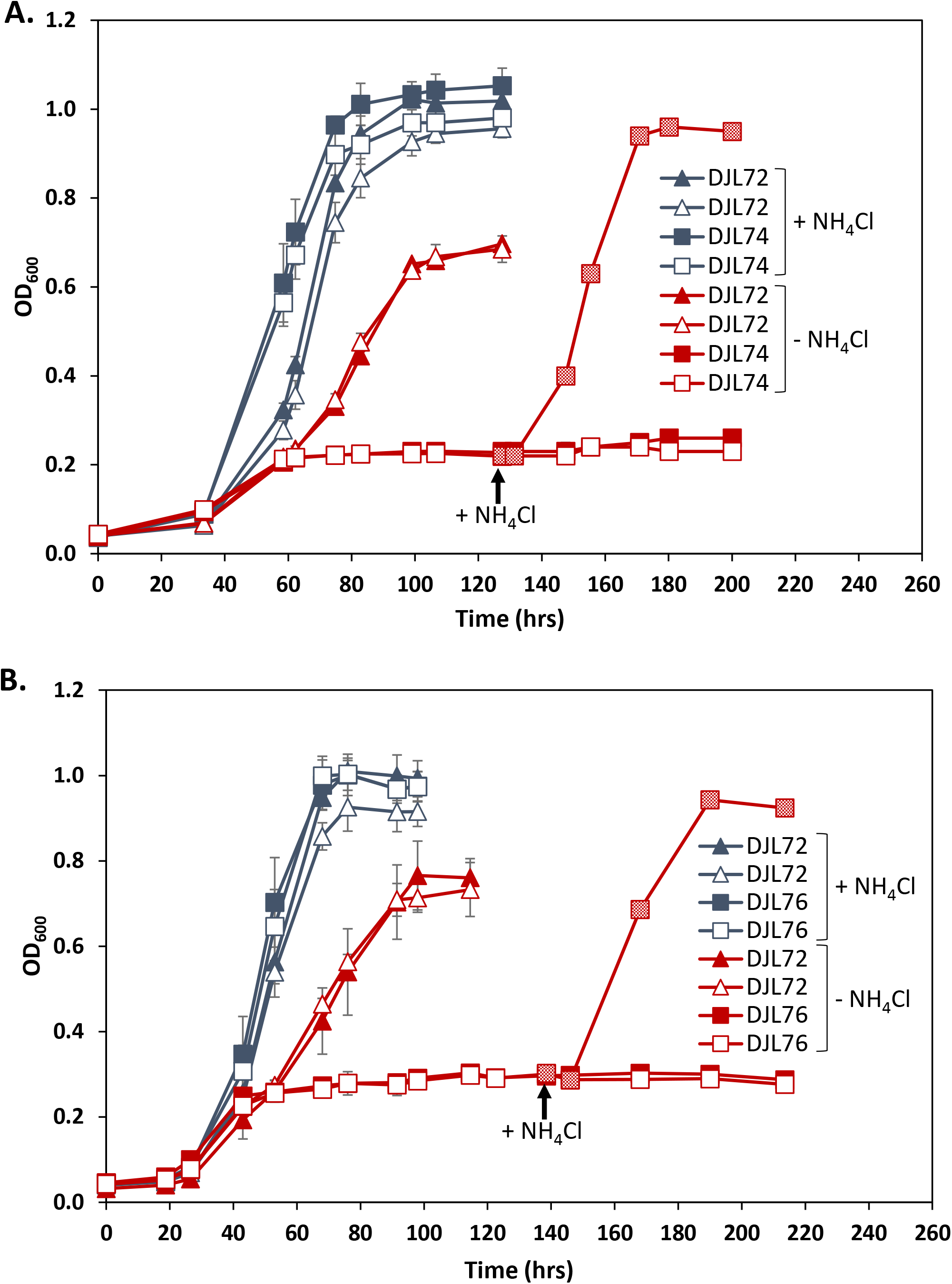
Comparison of the growth of *M. acetivorans* strain DJL72 to strains DJL74 (A) and DJL76 (B) with methanol +/− NH_4_Cl. Strains were grown in HS medium with 125 mM methanol and 18 mM NH_4_Cl as indicated. 50 μM tetracycline was added to cultures with open symbols. Data are the mean of four replicates and error bars represent standard deviation. Arrow denotes NH_4_Cl addition to cultures of DJL74 or DJL76.

qPCR analysis of the relative transcript abundance of *nifH* and *nifD* confirmed significant repression of the *nif* operon in strains DJL74 and DJL76, compared to strain DJL72 (**Fig. 5**). In both strains there was a >10-fold (>90%) reduction in both *nifH* and *nifD* transcript abundance in cells grown with or without NH_4_Cl. Thus, induction of the *nif* operon by the absence of NH_4_Cl is insufficient to overcome dCas9-mediated repression with either gRNA-*Pnif* or gRNA-*nifD*. Interestingly, although gRNA-*nifD* targets downstream of *nifH* (**Fig. 3**), significantly less *nifH* transcript was detected in cells of strain DJL76. It is possible blockage of RNAP by dCas9 bound at *nifD* results in abortive transcripts harboring *nifH* that are unstable and are subsequently degraded. Consistent with abolished diazotrophic growth, NifD was not detected by Western blot in lysate from cells of strains DJL74 and DJL76 (**Fig. 6**). These results reveal a 20-nt gRNA is enough to target dCas9 to block transcription of the *nif* operon preventing NifHDK production, and subsequently the ability to grow by fixing N_2_.

**Fig. 5.**
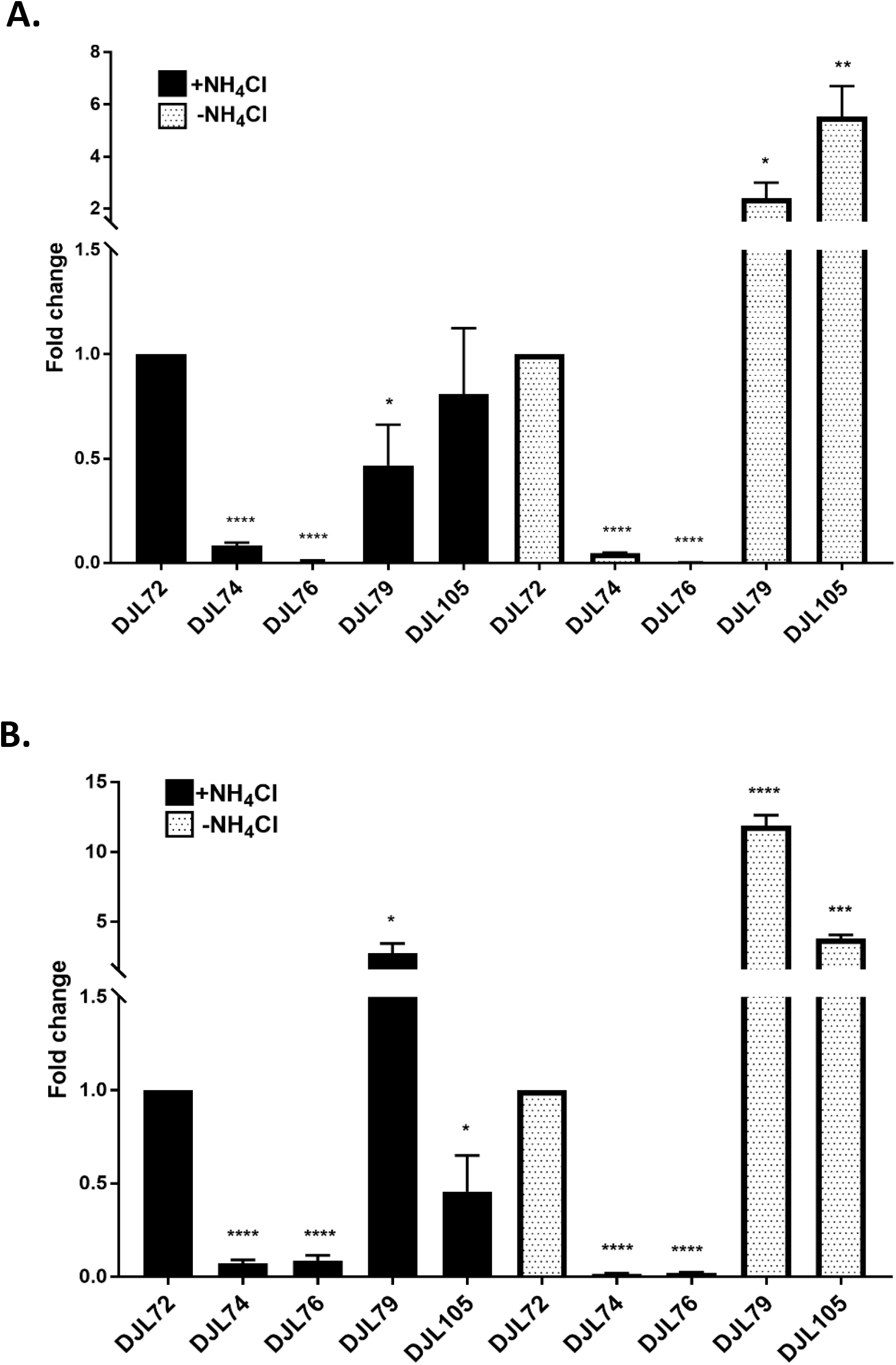
Relative transcript abundance of *nifH* (A) and *nifD* (B) in *M. acetivorans* CRISPRi-dCas9 strains. Strains were grown in HS medium with 125 mM methanol +/− NH_4_Cl. qPCR was performed as described in the material and methods. Data are the mean of three biological replicates analyzed in duplicate. The relative abundance of *nifH* and *nifD* in strain DJL72 (gRNA-less) was set to one to determine fold change in the other strains. *, **, ***, and **** denote p<0.05, < 0.01, p < 0.001, and p < 0.0001, respectively.

**Fig. 6.**
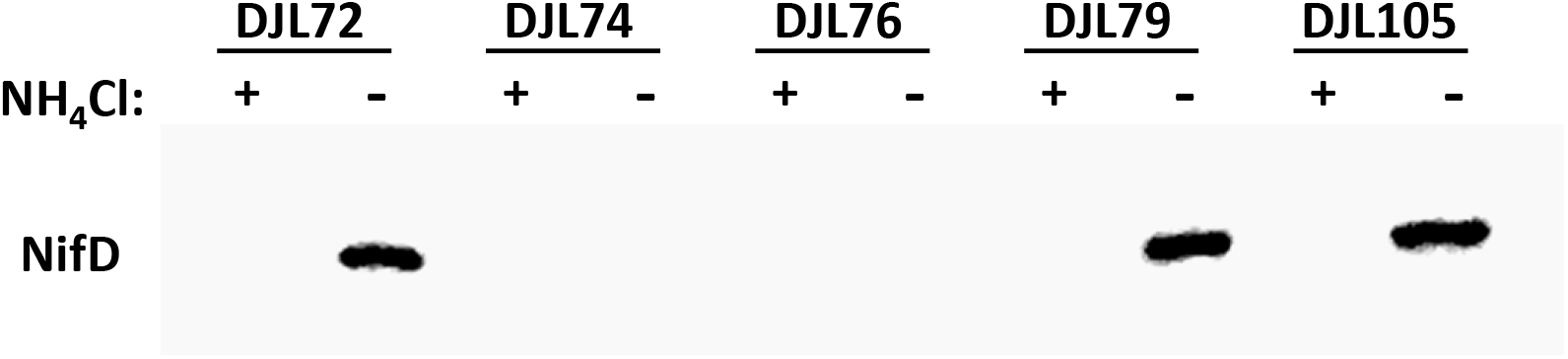
Comparison of NifD abundance in cell lysate from CRISPRi-dCas9 *M. acetivorans* strains. Strains were grown in HS medium with 125 mM methanol +/− NH_4_Cl. Western blot analysis was performed with antibodies specific to *M. acetivorans* NifD.

### Repression of *nifB* by dCas9 inhibits diazotrophic growth of *M. acetivorans*

To further ascertain the utility of the CRISPRi-dCas9 system to study gene function in *M. acetivorans*, we targeted the gene encoding the nitrogenase cofactor biogenesis protein NifB, the radical SAM-dependent enzyme required for the insertion of carbide into the M-cluster of Mo-nitrogenase [28]. Without the M-cluster, Mo-nitrogenase is inactive. Thus, NifB is essential for the ability of diazotrophs to fix N_2_. *M. acetivorans* contains a truncated version of NifB but biochemical evidence revealed it is capable of catalyzing steps leading to synthesis of the M-cluster [30]. However, the importance of NifB to diazotrophy by *M. acetivorans* has not been investigated. A gRNA (gRNA-*nifB*) was designed to target the sense strand starting ~100 nucleotides downstream of the start codon of *nifB* (**Fig. 3B**). *M. acetivorans* strain DJL105 was generated that harbors integrated pDL745 expressing gRNA-*nifB* (**Table 1**). Growth of strain DJL105 was identical to control strain DJL72 with methanol in HS medium containing NH_4_Cl (**Fig. 7A**). However, strain DJL105 exhibited delayed and slower growth in medium lacking NH_4_Cl. Analysis of *nifB* transcript abundance by qPCR revealed a 7- to 10-fold (>85%) reduction in strain DJL105 compared to strain DJL72 (**Fig. 8A**). Interestingly, the transcript abundance of *nifH* and *nifD* was 4 to 5-fold higher in cells of strain DJL105 compared to cells of strain DJL72 grown in the absence of NH_4_Cl (**Fig. 5**). However, the level of NifD in lysate from DJL105 cells was similar to the level of NifD in lysate from strain DJL72 cells as determined by Western Blot (**Fig. 6**). These results are consistent with NifB having a function specific to N_2_ fixation in *M. acetivorans*.

**Fig. 7.**
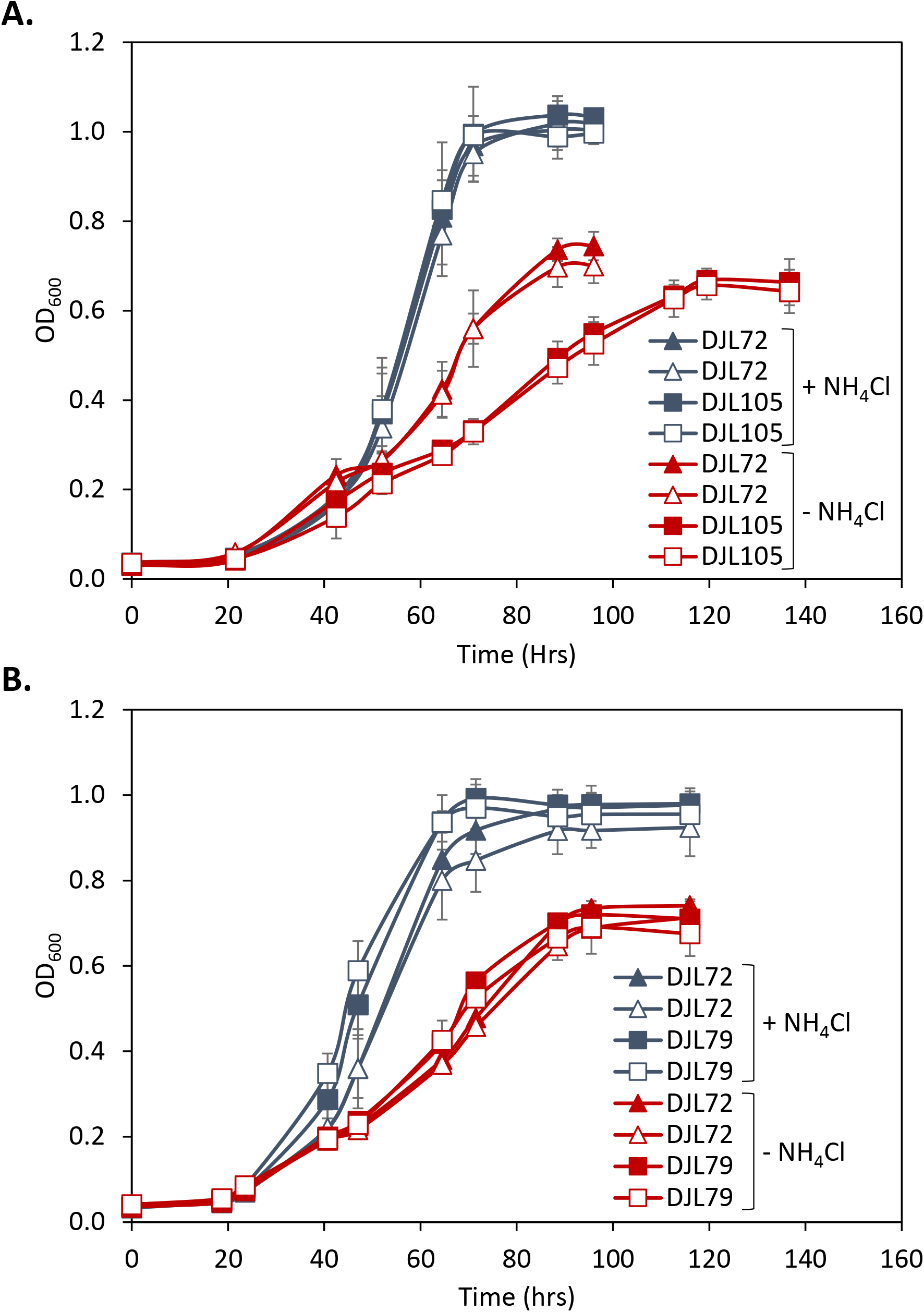
Comparison of the growth of *M. acetivorans* strain DJL72 to strains DJL79 (A) and DJL105 (B) with methanol +/− NH_4_Cl. Strains were grown in HS medium with 125 mM methanol and 18 mM NH_4_Cl as indicated. 50 μM tetracycline was added to cultures with open symbols. Data are the mean of four replicates and error bars represent standard deviation.

**Fig. 8.**
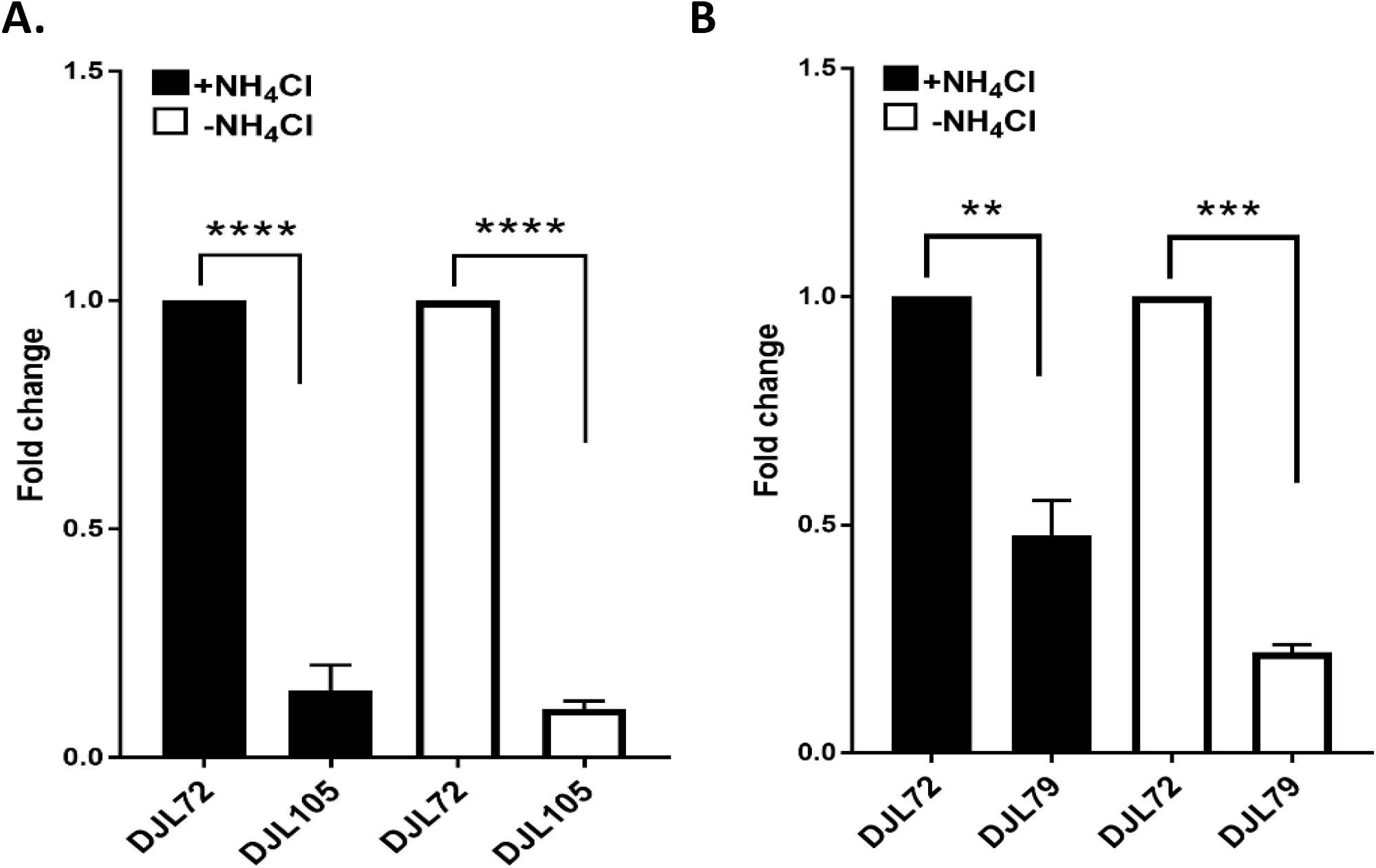
Relative transcript abundance of genes targeted for dCas9-mediated repression. **A)** Fold change in *nifB* transcript abundance in strain DJL72 versus DJL105. **B)** Fold change *nrpR1* transcript abundance in strain DJL72 versus DJL79. Strains were grown in HS medium with 125 mM methanol +/− NH_4_Cl. qPCR was performed as described in the material and methods. Data are the mean of three biological replicates analyzed in duplicate. **, ***, and **** denote p < 0.01, p < 0.001, and p < 0.0001, respectively.

### Repression of *nrpR1* by dCas9 affects *nif* operon expression

Finally, the ability to use the CRISPRi-dCas9 system to study genes involved in regulation of transcription in methanogens was tested. NrpR is the known repressor of *nif* operon transcription in methanogens [22, 31]. Although, *in vivo* analysis of NrpR has not been performed in *M. acetivorans*, the *nif* operon contains the identified NrpR operator (**Fig. 3A**) and recombinant *M. acetivorans* NrpR is functional in binding DNA [32]. Like other *Methanosarcina* species, NrpR is encoded by two separate genes in *M. acetivorans* [32]. NrpR1 contains the DNA-binding region, and deletion of only *nrpR1* in *Methanosarcina mazei* results in the loss of the *nif* operon repression [33]. Therefore, only *nrpR1* was targeted for dCas9-mediated repression in *M. acetivorans*. A gRNA (gRNA-*nrpR1*) was designed to target the sense strand starting six nucleotides downstream of the start codon of *nrpR1* (**Fig. 3C**). *M. acetivorans* strain DJL79 was generated with integrated pDL739 expressing gRNA-*nrpR1* (**Table 1**). Strain DJL79 and the control strain DJL72 exhibited identical growth in HS medium with or without NH_4_Cl (**Fig. 7B**). However, the transcript abundance of *nrpR1* was 2-fold (52%) and 5-fold (78%) reduced in DJL79 cells grown with or without NH_4_Cl, respectively (**Fig. 8**). Moreover, DJL79 had altered *nif* operon expression (**Fig. 5**). The transcript abundance of both *nifH* and *nifD* were significantly higher in DJL79 cells, compared to DJL72 control cells, grown in the absence of NH_4_Cl (*nif* inducing conditions). Cells of DJL79 grown in the presence of NH_4_Cl had more *nifD* detected, but less *nifH*, compared to the control cells. NifD was not detected in lysate from cells grown with NH_4_Cl (**Fig. 6**), despite the increased transcript abundance. The lower levels of *nifH* detected in strain DJL79 and the lack of NifD protein under *nif* repressing conditions (+NH_4_Cl) are consistent with significant post-transcriptional regulation of the *nif* operon [34–36]. Similar levels of NifD were observed in cells of strains DJL72 and DJL79 when grown under *nif* inducing conditions (−NH_4_Cl) (**Fig. 6**). These results demonstrate dCas9-mediated repression of NrpR1 alters transcription of the *nif* operon consistent with NrpR as the repressor of the *nif* operon.

## DISCUSSION

CRISPR-mediated transcriptional regulation is an extremely powerful tool for targeted and tunable control of gene expression. In bacteria such as *E. coli*, a single gRNA along with dCas9 is sufficient to elicit up to 99% repression of a synthetic target gene [10]. A gRNA targeting the sense (non-template) strand near the promoter or RBS effectively silences transcription initiation and elongation. However, using dCas9 with a single gRNA is less efficient in silencing transcription in eukaryotes (e.g. yeast) [37, 38]. Repression of target genes is improved by fusing additional regulatory domains to dCas9. An initial study showed fusing the mammalian repressor Mxi1 to dCas9 greatly enhanced targeted gene repression over dCas9 alone [39]. This difference may be due to the increased complexity of transcription initiation and elongation in eukaryotes. Although archaea have compact genomes, like bacteria, with genes lacking introns and arranged in operons, the archaeal transcription machinery is homologous to eukaryotic machinery (e.g. RNAP) and share similar mechanisms of transcription initiation and elongation [40]. However, like seen in bacteria, results in this study with *M. acetivorans* reveal expression of dCas9 with a single gRNA is sufficient to silence gene expression in archaea. Separate gRNAs targeting the sense strand of the *nif* operon promoter and near the start codon of *nifD*, repressed *nif* operon transcription, eliminated NifD production, and subsequently abolished growth with N_2_. The repression of the *nif* operon was efficient and stable; strains expressing either gRNA-*P_nif_* or gRNA-*nifD* were never able to overcome dCas9-mediated repression and grow with N_2_. Remarkably, *M. acetivorans* could not relieve repression to an extent that would allow production of sufficient nitrogenase to support growth, despite the strongest selective pressure (i.e. to grow). In fact, the cells remained in a viable but non-growing state for at least two weeks. The gRNAs targeting the promoter and the start codon worked equally well, indicating that dCas9 is capable of blocking transcription initiation and elongation in archaea. Importantly, based on the significant repression of genes targeted by gRNA-*nifD*, gRNA-*nifB*, and gRNA-*nrpR1*, in theory, any *M. acetivorans* gene could be silenced by expression of a single gRNA targeting the sense strand after the start codon.

dCas9-mediated repression of putative nitrogen fixation genes in *M. acetivorans* provides new insight, as well as confirms existing knowledge, about the expression, regulation, and maturation of nitrogenase in methanogens. These results confirm that the *nif* operon encodes Mo-nitrogenase, which is required for diazotrophic growth in medium containing molybdenum. *M. acetivorans* can also fix nitrogen using V- and/or Fe-nitrogenases in molybdenum-deplete medium (unpublished results). The regulation of the alternative nitrogenases has not been investigated in methanogens. Interestingly, *M. acetivorans* did not compensate for the loss of Mo-nitrogenase by producing an alternative nitrogenase to fix N_2_, at least under the conditions tested, which has been seen in some diazotrophic bacteria [41]. These results indicate that molybdenum is likely the critical effector molecule that controls expression of V- and Fe-nitrogenases in *M. acetivorans*.

NifB is essential for the activity of all nitrogenases [42]. However, dCas9-mediated repression of *nifB* did not abolish diazotrophic growth of strain DJL105, unlike the repression of the *nif* operon. NifB converts [4Fe-4S] clusters into NifB-co, an [8Fe-9S-C] cluster that is the precursor to the active site cluster (M-cluster; [7Fe-9S-C-Mo-homocitrate]) in NifDK [28]. The genome of *M. acetivorans* encodes a single NifB that lacks a NifX-like domain found in some bacteria. Recombinant *M. acetivorans* NifB supports the *in vitro* synthesis of the M-cluster in NifDK [30]. Unlike the structural components (NifDK) of Mo-nitrogenase which are needed in high abundance [43], only a small amount of NifB is likely required for maturation of nitrogenase. Therefore, the impaired growth of strain DJL105 in the absence of NH_4_Cl could be due to minimal production of NifB. Alternatively, an unknown protein could compensate for the loss of NifB production. Additional experimentation is required to confirm that NifB encoded by MA4195 is solely required for the maturation of nitrogenase. Interestingly, a significant increase in the transcript abundance of *nifH* and *nifD* was observed in strain DJL105 cells grown without NH_4_Cl (**Fig. 5**). NifB limitation likely decreases the level of functional Mo-nitrogenase. To compensate, the cells increase *nif* operon transcription to attempt to produce more Mo-nitrogenase. These results provide new insight into the relationship between nitrogenase maturation and expression in methanogens.

NrpR is a transcriptional repressor of the nitrogen fixation and assimilation genes in methanogens and has been extensively studied in *M. maripaludis and M. mazei*, but not *M. acetivorans* [22, 31–33, 44]. *M. acetivorans* strain DJL79 had higher transcript abundance under nitrogen sufficient conditions (+NH_4_Cl) compared to the control, revealing NrpR is also the repressor of the *nif* operon in this methanogen. Increased transcription of the *nif* operon in strain DJL79 did not result in increased NifD production, consistent with additional post-transcriptional regulation, and underscores the complexity of nitrogenase regulation in diazotrophs [34, 36]. Interestingly, *nifH* and *nifD* transcript abundance was highest in strain DJL79 under nitrogen limitation, indicating NrpR may primarily function to titer the level of transcription during nitrogen limitation (−NH_4_Cl).

Compared to the endogenous-based CRISPRi systems in *Haloferax* and *Sulfolobus* for silencing genes in archaea, the exogenous dCas9-based CRISPRi system in *Methanosarcina* is simpler and more efficient. The *Haloferax* CRISPRi system requires the use of mutant strains deleted of all or parts of the endogenous type I CRISPR components [18, 19]. Thus, use of the system is restricted to specialized strains and since the endogenous system is used, it is unclear if these mutations cause unknown effects. Moreover, the *Haloferax* system only blocks transcription initiation, not elongation, because only gRNAs that target the template strand near the transcriptional start site result in significant repression [18, 19]. In contrast to the *Methanosarcina* CRISPRi-dCas9 and *Haloferax* CRISPRi systems, the *Sulfolobus* CRISPRi system degrades existing mRNA using endogenous type III CRISPR system components. Greater than 90% reduction in mRNA and subsequent protein activity can be achieved by the *Sulfolobus* system [17]. However, optimal mRNA reduction requires multiple gRNAs and/or increased gRNA dosage [15, 17]. In contrast, the *M. acetivorans* CRISPRi-dCas9 system requires only two components, dCas9 and a single gRNA to efficiently block both transcription initiation and elongation. Although specialized strains of *M. acetivorans* are needed to integrate pDL730 and derivatives into the chromosome, these strains are not required if the plasmid is first converted into the replicating version (**Fig. 1**). Therefore, any existing *M. acetivorans* strain could be transformed with replicating pDL730 and derivatives allowing targeted gene repression in existing mutants. Although not demonstrated here, we expect that expression of multiple gRNAs targeting different genes/operons in a single strain will result in simultaneous repression, and we are currently testing this possibility.

There are limitations to the current CRISPRi-dCas9 system. Like the *Haloferax* and *Sulfolobus* CRISPRi systems, there is a lack of control. It is not possible to turn on/off repression of a target gene/operon. Like the *M. acetivorans* CRISPR-Cas9 system, where tetracycline was not required to induce genome editing [24], tetracycline had no effect on dCas9-mediated repression of target genes in the CRISPRi-dCas9 strains. This limits the ability to study essential or critical genes since the system is stuck in the “on” state. We are currently attempting to engineer additional control in the system.

In summary, the results in this study demonstrate that the CRISPRi-dCas9 system is a simple, yet powerful, new tool to understand and control the biology of *M. acetivorans*. Importantly, since no additional processing of transformants is required, CRISPRi-dCas9 strains can be generated in as little as two weeks (including plasmid construction). The system will expedite understanding gene function, regulation, and interaction in methanogens, and can likely be adapted for use in other non-extremophilic archaea as well.

## Acknowledgments

We thank Dipti Nayak and Bill Metcalf for providing *M. acetivorans* strains and plasmids. This work was supported in part by DOE Biosciences grant number DE-SC0019226 (DJL), NSF grant number MCB1817819 (DJL), and the Arkansas Biosciences Institute (DJL), the major research component of the Arkansas Tobacco Settlement Proceeds Act of 2000.

## MATERIALS AND METHODS

### *M. acetivorans* strains and growth

*M. acetivorans* strains used are listed in **Table 1**. *M. acetivorans* strain WWM73 served as the parent strain for all experiments. HS medium was prepared as previously described [45], except NH_4_Cl was omitted. Methanol, sulfide, tetracycline, puromycin, and NH_4_Cl were added from anaerobic sterile stocks using sterile syringes prior to inoculation. All *M. acetivorans* strains were grown at 35 °C in Balch tubes containing 10 ml HS medium with 125 mM methanol and 0.025% sulfide. Headspace gas composition was 75% N_2_, 20% CO_2_, and 5% H_2_. Tetracycline (50 μM), puromycin (2 μg/ml), and NH_4_Cl (18 mM) were added where indicated. Growth was measured by monitoring optical density at 600 nm (OD_600_) using a spectrophotometer.

### Construction of *M. acetivorans* CRISPRi-dCas9 plasmids and strains

All primers and synthetic DNA fragments (gblocks) listed in **Tables S1** and **S2** were designed using Geneious Prime and purchased from Integrated DNA Technologies (IDT). To construct a universal CRISPRi-dCas9 plasmid, *cas9* was replaced with *dcas9* in pDN203 (gift from Metcalf lab) [24]. First, dCas9 with PCR amplified using Q5® High-Fidelity DNA Polymerase (New England Biolabs) with specific primers (Table S1) and pMJ841 as a template. The *dcas9* PCR amplicon was purified using a Wizard SV Gel and PCR Clean-up kit (Promega) and digested with *Asc*I and *Hind*III. pDN203 was similarly digested with *Asc*I and *Hind*III and the plasmid lacking *cas9* was purified from an agarose gel using the Wizard SV Gel and PCR Clean-up kit. Digested *dCas9* was ligated into pDN203 using T4 DNA ligase followed by transformation of *E. coli* DH5α competent cells (New England Biolabs). Transformants were screened by PCR and the final universal CRISPRi-dCas9 plasmid (pDL730) containing *dcas9* was verified by sequencing (Eurofins Genomics LLC, USA). All other plasmids to target specific genes and operons (**Table S3**) for dCas9-mediated repression were constructed by introducing a gblock that contains a specific gRNA into *Asc*I-digested pDL730 using Gibson Assembly® Ultra Kit (Cat# GA1200-10, Synthetic Genomics, Inc.) according to the manufacturer’s instructions. CRISPRi-dCas9 plasmids were converted into *M. acetivorans* replicating plasmids by retrofitting with pAMG40 [46] Gateway BP Clonase II kit (Invitrogen) according to manufacturer’s instructions. *M. acetivorans* strain WWM73 was separately transformed with replicating and non-replicating plasmids using a liposome-mediated transformation protocol as described [47]. Transformants were selected by anaerobic growth on agar plates containing 125 mM methanol and 2 μg/ml puromycin. Colonies were screened by PCR and a positive colony was selected as the new CRISPRi-dCas9 strain (**Table 1**). Strains were maintained in HS medium with 125 mM methanol and 2 μg/ml puromycin.

### Gene expression analysis

*M. acetivorans* cells (4 ml of culture) were harvested at mid-log phase (OD_600_ = 0.2 to 0.4) by centrifugation at 5,000 × *g* inside an anaerobic chamber (Coy Laboratories). The cell pellets were resuspended in 1ml Trizol (Life Technologies, Cat #15596-026) and stored at –80 °C until use. Total RNA was purified from cells using a Direct-zol RNA Miniprep kit (Zymo research, Cat# R2050) according to the manufacturer’s instructions followed by an additional DNase treatment using DNA-free™ DNA Removal Kit (Thermo Fisher Scientific, Cat# AM1906). RNA concentrations were determined using a Thermo Scientific™ NanoDrop 2000. To determine transcript abundance by quantitative real-time PCR assay (qPCR), cDNA was generated with 300ng of total RNA using iScript™ Select cDNA Synthesis Kit (Bio-Rad, Cat# 1708897) with the following conditions (25 °C for 5 min, 42 °C for 30 min, and 85 °C for 5 min). cDNA was stored at −20 °C until use. Each qPCR reaction (10 μl) contained 1x SsoAdvaced universal SYBR Green Supermix (Bio-Rad, Cat#1725271), 300 nM of forward and reverse primers (**Table S1**), and 300-fold dilution cDNA. The qPCR reactions were performed using a CFX96 Real-time PCR detection system (Bio-Rad) for all samples under the following conditions: 95 °C for 1 min followed by 37 cycles of (95 °C for 20 s, 60 °C for 20s) followed by melting curve analysis to ensure the specificity of amplification. qPCR data were analyzed using ΔΔcq method (R) with 16sRNA used as an internal control. To determine the transcript abundance of a specific gene, Δcq values of the gene in the samples were calibrated to the average of its Δcq values in the control group (DJL72 cells) unless indicated otherwise. Duplicate qPCR reactions were conducted for three biological replicates for each strain.

### Western blot analysis

*M. acetivorans* cells were harvested at mid-log phase (OD_600_ = 0.2 to 0.4) by centrifugation at 5,000 × *g* inside an anaerobic chamber (Coy Laboratories). Cell pellets were resuspended in buffer A (50 mM Tris-HCl, pH 8.0, 150 mM NaCl, 1 %, Tween 20, 1 mM Benzamidine hydrochloride hydrate, and 2 mM EDTA), sonicated 2-3 times in ice bath using a Branson Digital Sonifier 450 (Branson Ultrasonics) and centrifuged at 14,500 × *g* for 3 min. The total protein concentration in the cell lysates were measured by the Bradford method using Bio-Rad Protein Assay Dye Reagent Concentrate (Cat #5000006) according to the manufacturer’s instructions. Protein (5 μg) samples were resolved in a 10 % SDS-PAGE gel and transferred to an Amersham™ Protran™ NC membrane (Fisher Scientific, Cat#10600005). The membrane was blocked with blocking buffer (5 % skim milk in Tris-Buffered Saline with 0.1% Tween, TBST) for 15 min at room temperature (RT), and probed with a custom anti-NifD antibody (GenScript) overnight. The membrane was washed at least three times by TBST for 5min each and then incubated with the secondary antibody (goat anti-rabbit) for 1 h at RT with slow shaking. After three washes, the membrane was developed with an ECL reagent (Amersham ECL™ Prime Western Blotting detection reagent from GE Healthcare UK, Cat# RPN2232) and scanned by FluorChem™ 8900 (Alpha Innotech, San Leandro, CA). Western blot analysis of lysate from all strains were done with at least three biological replicates.

